# Upregulation of photosynthetic capacity and thylakoid membrane protein enhanced tolerance to heat stress in wucai (*Brassica campestris L*.)

**DOI:** 10.1101/493247

**Authors:** Yujie Yuan, Lingyun Yuan, Jie Wang, Mengru Zhao, Yun Dai, Shilei Xie, Shidong Zhu, Jingfeng Hou, Guohu Chen, Chenggang Wang

## Abstract

The hot climate of southern China from late summer to early fall is one of the major factors limiting the yield and quality of wucai (*Brassica campestris L*.). Under high temperature stress, heat-tolerant cultivars presented moderate injury to the photosynthesic apparatus, less inhibition of photochemical activity, better osmotic adjustment and antioxidant defences capacity compared to heat-sensitive cultivars. To study the effects of high temperature on the growth and development of wucai, plants of WS-1 (heat-tolerant) and WS-6 (heat-sensitive) were exposed to four heat stress treatments in growth chambers for 3 days. Chloroplasts of two cultivars evaluated for photosynthetic characteristics, fatty acid composition and differentially expressed proteins of thylakoid membrane. The chlorophyll (Chl) content was markedly reduced by heat stress, inhibiting photochemical activity. However, larger decreases in growth and photosynthetic parameters [net photosynthetic rate (P_N_), stomatal conductance (G_S_), intercellular CO_2_ concentration (C_i_), and leaf transpiration rate (E)] occurred under heat stress in WS-6, compared with WS-1. In addition, WS-6 showed an obviouse K point in O-J-I-P steps under extremely high temperature, which indicated OEC had been damaged. WS-1 showed higher of maximum quantum yield of primary PSII photochemistry (F_V_/F_M_), number of active reaction centres per cross section of PSII (RC/CS_M_), average absorbed photon flux per cross section of PSII (ABS/CS_M_), maximum trapped exciton flux per cross section of PSII (TR_0_/CS_M_), electron transport flux from Q_A_ to Q_B_ per cross section of PSII (ET_0_/ CS_M_) and performance index on absorption basis (PIABS) which indicated greater heat stability in terms of PSII function under higher temperature. Compared to WS-6, WS-1 showed higher membrane stability and photochemical efficiency, and greater increase of saturated fatty acids (SFA), especially palmitic acid under heat stress. WS-1 had higher recovery rate compared to WS-6 after 41°C heat stress treatment. Additionally, two-dimensional blue native/SDS-PAGE analysis of chloroplast was carried out to compare the differentially expressed proteins between two cultivars. We obtained seven major protein complexes included supercomplexes, PSI-LHCII/PSII monomer, PSII monomer, CP43 less PSII/ATP synthase, LHCII trimer, LHCII monomer and ATP synthase after first dimentional seperation in both cultivars after the first dimensional separation. Then ten differential membrane proteins included light-harvesting Chl a/b-binding (LHC) protein, ATP synthase subunit alpha, ATP synthase subunit beta, photosystem I P700 chlorophyll a apoprotein A2, photosystem II CP43 reaction center protein, photosystem II D2 protein and photosystem II

OS have been found between WS-1 and WS-6. These differentially proteins in cellular membranes could contribute to the differential level of heat tolerance between two wucai cultivars. Our results demonstrated that the heat-tolerant cultivar WS-1 had a greater capacity for photosynthesis and membrane stability by upregulating proteins abundance including light harvesting (light-harvesting Chl a/b-binding protein), energy metabolism (ATPase), and proteins of PSII reaction center (D2, CP43) under heat stress.

## Introduction

Wucai (*Brassica campestris L*. ssp. chinensis var. rosularis Tsen et Lee.) is a type of non-heading Chinese cabbage with high nutritional value. It is a cruciferous, biennial herb, originated from China and distributed mainly in Yangtze-Huaihe River basin (Yuan and Sun 2001). Wucai grows well in cold weather of late October, but not in the hot summer (Shao et al. 2014). The high temperature might inhibit the seedling growth of wucai and even cause heat damage. To achieve annual production and meet market demand, it is critical to select and breed heat-tolerant wucai cultivars.

Photosystem II is generally considered to be the primary target of heat-induced inactivation of the photosynthetic membranes (Allakhverdiev et al. 2008; Mohanty et al. 2012). Environmental stress mediated decreases in photosynthesis may result from inhibition of PSII activity, which also leads to a decrease in variable chlorophyll fluorescence (Baker and Rosenqvist 2004; Maxwell and Johnson 2000; Feng et al. 2014). Chlorophyll *a* fluorescence is widely used in photosynthesis research, plant physiology, plant phenotyping, remote sensing of plants, and other fields of research that are related to photosynthesis (Kalaji et al. 2017, Mishra et al. 2016). Specifically, the analysis of fluorescence signals provides detailed information on the status and function of Photosystem II (PSII) reaction centers, lightharvesting antenna complexes, and both the donor and acceptor sides of PSII (Kalaji, H et al. 2016).

The membrane plays important roles in sensing environmental change, signal transduction and substance metabolism (Mittler et al. 2012; Guyot et al. 2015). Phospholipids forms the bilayers of the membrane, which mainly consists of fatty acids in saturated or unsaturated forms, and proper fatty acid composition is critical for maintaining membrane stability during plant adaptation to stress conditions (Gigon et al. 2004). Up-shifts in temperature increase the fluidity of the cytoplasmic membrane and causes reduction of unsaturated fatty acid content and increase in saturated fatty acid content, leading to increased saturation level of fatty acids, which has been positively associated with heat tolerance (Larkindale and Huang, 2004).

Denaturation or dissociation of membrane proteins related to photosystem II, including 33-kDa manganese (Mn)-stabilizing protein (Yamane et al., 1998), oxygen evolving complex (OEC) (De Ronde et al., 2004) and D1, D2 proteins of the reaction center (Yamamoto Y. et al. 2008) has been reported under heat stress. Sevral studies roported a lesser or later decrease of membrane proteins was observed in a heat tolerant line of bentgrass (Agrostis spp.) compared to a heat sensitive line, including those categorized to energy metabolism (ATP-synthase, Cytochrome b6f, chloroplast oxygen-evolving enhancer protein, and pyruvate dehydrogenase kinase) and antioxidant processes (catalase and peroxidase) in response to heat stress (Jespersen et al., 2015).

Here we used this approach using blue-native PAGE (BN-PAGE), which substantially improved recovery of dechlorinating activity after electrophoresis, resulting in higher sensitivity and enabling analysis of a wider range of substrates. The blue-native PAGE is widely used in membrane protein complexes, and it has been uesd successfully ito characterize respiratory complexes in yeast mitochindria (Cruciat et al. 2000), photosythetic complexes in plant (Ciambella et al. 2005), and cell cell envelope protein complexes in *E.coli* (Pan et al. 2010). Moreover, recent modifications have made it possible to apply this method to the study of whole protein complexes of an organism (Jha et al. 2016). The two-dimensional blue native/sodium dodecyl sulfate polyacrylamide gel electrophoresis (2D BN/SDS-PAGE) is a method to investigate protein complexes (Lasserre et al. 2012).

In our previous study, it was suggested that heat stress could reduce Chl content that inhibited photochemical activity and caused sharply decrease in growth and photosythetic parameters. Additionally, we found heat-sensitive cultivars had a greater accumulation of reactive oxygen species (ROS) and malondialdehyde (MDA) that caused greater severity of damage to the photosynthetic apparatus and membrane system relative to heat-tolerant cultivars. We believed that heat-tolerant cultivars had greater a greater capacity for maintaining leaf RWC and scavenging ROS was due to better osmotic adjustment and antioxidant defences capacity, as compared with WS-6 (Zou et al., 2016 and 2017). Although photosythetic parameters, osmotic adjustment and antioxidant defence have been well documented, limited information is available on changes in membrane proteins for plant adaptation to heat stress, particularly on specific membrane proteins that could be changed conferring membrane thermostability and plant tolerance to heat stress.

The present study was carried out with two wucai cultivars (heat tolerant and heat sensitive) to investigate photosynthetic capacity and membrane fatty acid composition. We isolated intact chloroplasts and obtained thylakoid membranes, and analysed membrane protein complexes and diffrential proteins under heat stress. The objective of this study was to assess comparative adaptation changes responding to heat stress and to explore the regulation mechanism from fatty acid composition and differential expression proteins of heat-tolerant cultivar, which could provide a guidance and theoretical basis for wucai tolerant breeding.

## Materials and methods

### Plant cultivation and treatments

Wucai seeds of WS-1 (heat-tolerant) and WS-6 (heat-sensitive) were supplied by the Vegetable Genetics and Breeding Laboratory at Anhui Agricultural University, China. Young seedlings were planted in plastic pots filled with sterilized soil and grown at 20/12 °C day/night) with a 15h photoperiod, a photon flux density (PFD) range of 300–500μmol m^−2^ s^−1^, and a relative humidity range of 60–70% in a green house. When the sixth leaf was fully expanded (approximately 4 weeks old), the uniform size seedlings were adapted in an illuminated incubation chamber (GXZ-260C) for 2 d before treatment. Seedlings were randomly separated into four groups for heat stress treatment. The day/night (d/n) temperatures of the five treatments were as follows:

a. Cont, 20°C/12°C (d/n) treatment for 3 days
b. TS, 27°C/18°C (d/n) stress treatment for 3 days
c. TF, 34°C/24°C (d/n) stress treatment for 3 days
d. FO, 41°C/30°C (d/n) stress treatment for 3 days
e. After 41°C/30°C (d/n) stress treatment for 3 days, transferred to the control level for 3 days.

### Morphological parameters

Morphological parameters include plant height, stem diameter, leaf length, leaf width and single plant weight. Morphological parameters were obtained from ten seedlings in each replication. Plant height was measured from cotyledonary node to the top with a ruler. Stem diameter was measured at the cotyledonary node using a vernier caliper. After the measurements, all seedlings were cut at the bases of their stems and rinsed thoroughly with distilled water, and single plant weight was measured after removal of residual moisture. Leaf area and RWC were measured from 10 seedlings in each replication. Data were averaged from three replicates. The area of the 3rd expanded leaf (from the core) was measured with an Expression 1680 scanner (Epson, Sydney, Austrialia). Leaf RWC was measured according to the method of Barrs and Weatherley (1962). After fresh weight (FW) was measured, the leaves were floated on deionised water for 6h under low irradiance. Then the leaves were blotted to wipe off excess water, weighed to record fully turgid weight (TW), and then subjected to an oven drying at 75°C for 72h to record the dry weight (DW). Leaf RWC was calculated by the following formula: Leaf RWC(%) = (FW-DW)/(TW-DW)×100%.

### Chlorophyll content

The Chl contents were measured as described by Strain and Svec (1966) with some modifications. Fresh leaf samples (0.2 g) were obtained from the fragments of the third leaves in each treatment group and incubated for 24 h in the dark at 4 °C in 25 mL of acetone and ethanol and water at 4.5:4.5:1 (v/v/v); after filtration, absorbance values were then recorded at 649 and 665 nm. The Chl concentrations were calculated using the following formulae: Chl a = 13.95 A665 - 6.88 A649; Chl b = 24.96 A649 - 7.32 A665; and Chl a+b = Chl a + Chl b.

### Photosynthetic parameters

The gas exchange parameters evaluated were P_N_, G_S_, C_i_, and E. Measurements were taken with a portable photosynthesis system (LI-6400, LI-COR Inc., Lincoln, NE, USA) on fully expanded third leaf. Ten replicate measurements were taken per treatment on a clear day between 09:00 am and 11:00 am. The conditions during measurements were 25 °C, RH of 70%, external CO_2_ concentration of 380 ± 10 lmol mol^−1^, and light intensity of 1,000 μmol photons m^−2^ s^−1^.

### Chlorophyll fluorescence parameters

The Chl fluorescence parameters were measured using a handheld portable fluorometer (Pocket PEA). Fully expanded third leaf was dark-adapted for 30 min prior to measurement. The different steps of the OJIP transient determined with the Photon Systems Instruments include fluorescence intensity at 50 ms considered as the initial fluorescence value, F_0_; maximum fluorescence level of the OJIP transient (FM) measured under saturating light conditions while intermediate fluorescence values were measured at 300 ms, 2 ms, and 60 ms, and labelled as F_300ms_, Fj and Fi respectively (Strasser et al. 2000). The OJIP transient was analysed with the JIP-test (Strasser et al. 2000, 2004). The test provides several parameters characterising the photosynthetic samples including estimates of energy fluxes per active reaction centre, and estimates of the probability/efficiencies of energy flow in the PSII (Strasser et al. 2000, 2004).

### Fatty acid composition of thylakoid membranes

Thylakoid membrane lipids were separated from the isolated thylakoid membranes and the fatty acids were analysed according to the method of (Zhang et al. 2010). The lipids extracts were separated by gas chromatography (HP6890, Agilent). An Agilent HP-INNOWax column (33m× 0.25mm) was packed with polyethylene glycol. Hydrogen flame ionization was detected at 230 °C, and the column temperature was programmed to rise from 170 °C to 210 °C at 5 °C per min. Chromatograms were recorded and peak areas were calculated to measure the fatty acids levels. Peaks were identified by comparisons against several external qualitative standards.

### Isolation and fractionation of chloroplast

Intact chloroplasts from the third fully expanded leaves were isolated and purified on percoll gradients according to the method of (Shu et al. 2015). To obtain the thylakoid membranes, the intact chloroplasts were ruptured in 50mM Hepes-KOH (pH 7.6) and 2mM MgCl_2_ at 4°C and the thylakoid membranes were collected by centrifugation at 14, 000g for 15min.

### Solubilization of thylakoid membrane proteins

Thylakoid membranes were resuspended in solubilisation buffer containing 25 mM BisTris-HCl (pH 7.0), 20% glycerol and 2% n-dodecyl-β-D-maltoside (Sigma). After incubation for 30 min on ice, samples were centrifuged at 14, 000g for 3 min to remove insoluble material. The supernatant was supplemented with a certain volume of sample buffer [1% (w/v) Coomassie brilliant blue G-250, 0.1 M BisTris-HCl, pH 7.0, 30% sucrose and 0.5 M 6-amino-n-caproic acid]. Dye-labelled protein samples were directly loaded onto Blue-native gels.

### Two-dimensional Blue-native/SDS-polyacrylamide gel electrophoresis

Two-dimensional Blue-native/SDS-polyacrylamide gel electrophoresis (2D BN/SDS-PAGE) was carried out as described by Reisinger & Eichacker49 with minor modifications. The first dimension, BN-PAGE, is a native polyacrylamide gel in which a gradient gel of 5%–13.5% acrylamide was used. The anode buffer contained 50 mM BisTris-HCl, pH 7.0 and the cathode buffer contained 50 mM Tricine, 15 mM BisTris and 0.01% (w/v) Coomassie brilliant blue G-250. The gel was run at 4 °C. When the BN-PAGE was completed, the gel was equilibrated in 6 M urea, 5% (w/v) SDS, 10% β-mercaptoethanol, 20% (v/v) glycerol, and 50 mM Tris-HCl (pH 7.0) for 20 min. After washing with deionized water for three times, individual lanes were cut and inserted into the gel loading hole of the second dimension, SDS-PAGE, which was carried out as described by Laemmli50. Protein spots were visualized using Coomassie brilliant blue R-250.

### Image acquisition and data analysis

The stained gels were scanned using the Image scanner III (GE Healthcare). The images were analyzed with ImagemasterTM 2D Platinum software version 6.0 (GE Healthcare). Three gels for each treatment from three independent experiments were used for the analysis. The intensities of spots were quantified based on the ratio of the volume of a single spot to the whole set of spots. Only spots with quantitative changes of at least 1.5-fold in abundance that were reproducible in three replicates were used for mass spectrometry.

### Statistical analysis

All data were statistically analyzed with SAS 13.0 software (SAS Institute, Inc., Cary, NC, USA) using Duncan’s multiple range test at a 0.05 level of significance.

## Results

### Morphological parameters analyses

With increasing temperature, 27°C/18°C controls slightly influence in fresh weight, dry weight, plant height, stem diameter, leaf width and leaf length. In contrast, 34°C/24°C controls and 41°C/30°C controls significantly inhibited the growth of plants. Fresh weight, dry weight, plant height and stem diameter of both cultivars were decreased gradually showing decreases of 9.58%, 12.73%, 8.07%, 7.77% and 16.11%, 16.50%, 16.23%, 20.76% in WS-1 and WS-6 at 40°C, respectively, compared to 20°C controls (Table 1). The leaf width and leaf length were sharply decreased of 5.80%, 3.53% and 9.19%, 6.84%. Moreover, larger increases in fresh weight, dry weight, plant height, stem diameter leaf width and leaf length occurred at 35°C and 40°C in WS-6, compared with WS-1.

**Table 1.** Effects of heat stress on fresh weight, dry weight, plant height, stem diameter, leaf width and leaf length in wucai plants

| Controls |  | Fresh weight | Dry weight | Plant height | Stem diameter | Leaf width | Leaf length |
| --- | --- | --- | --- | --- | --- | --- | --- |
| WS-1 | Cont | 30.80±0.149a | 2.92±0.038a | 10.27±0.25c | 3.68±0.011a | 4.83±0.15a | 6.24±0.11a |
|  | TS | 30.76±0.118a | 2.91±0.026a | 10.32±0.16c | 3.65±0.012a | 4.84±0.15a | 6.26±0.14a |
|  | TF | 29.97±0.092b | 2.76±0.026b | 10.87±0.20b | 3.56±0.023b | 4.74±0.13a | 6.13±0.15ab |
|  | FO | 27.85±0.141c | 2.55±0.029c | 11.32±0.16a | 3.39±0.024c | 4.55±0.19b | 6.02±0.25b |
| WS-6 | Cont | 28.67±0.146a | 2.71±0.028a | 10.46±0.17c | 3.16±0.022a | 4.46±0.17b | 7.46±0.13a |
|  | TS | 28.63±0.118a | 2.69±0.023a | 10.48±0.14c | 3.13±0.037a | 4.48±0.18a | 7.48±0.07a |
|  | TF | 27.07±0.081b | 2.50±0.031b | 11.32±0.21b | 2.89±0.062b | 4.32±0.17c | 7.24±0.12b |
|  | FO | 24.05±0.081c | 2.26±0.024c | 12.30±0.19a | 2.50±0.048c | 4.05±0.18d | 6.95±0.14c |

### Leaf RWC

The general trend in leaf RWC response to heat stress was similar between both cultivars, while it differed in the magnitude of change (Fig 2). At 27°C, leaf RWC was not affected in either of the cultivars. However, it was decreased by 4.52% and 8.22% under 35°C and 40°C stress conditions in WS-1, while the decrease was by 11.95% at 34°C and 16.15% at 41°C in WS-1, respectively. After plants were transferred to moderate temperature condition for 3 days, average recovery rates were 98.29% of WS-1 and below 87.90% of WS-6. While the recovery of leaf RWC from 41°C heat stress, WS-1 occurred more quickly and mostly reached that of the control level.

**Fig.1.**
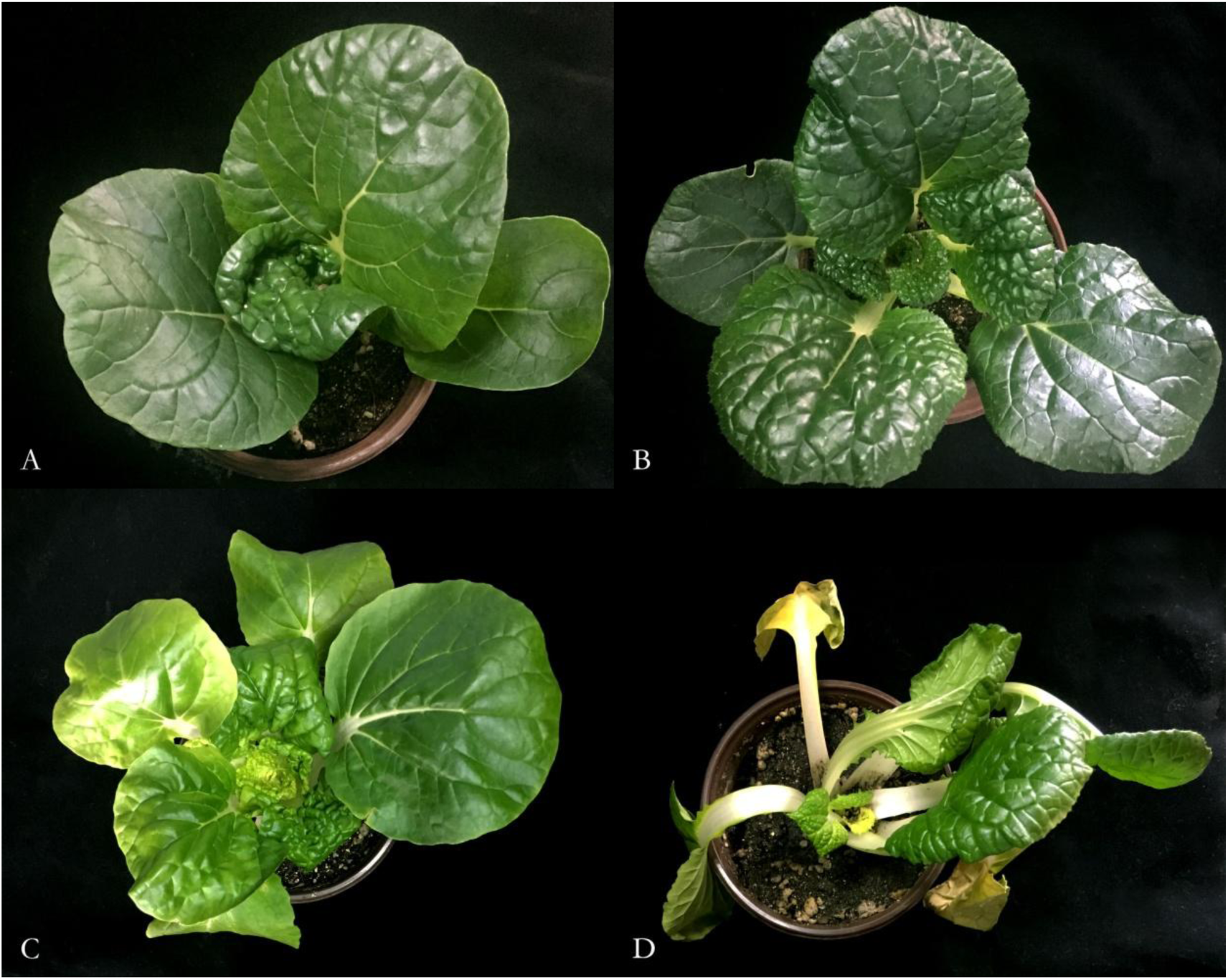
Performance in wucai plants. A: WS-1 *Cont* treatment, B: WS-6 *Cont* treatment, C: WS-1 *FO* treatment, D: WS-6 *FO* treatment. *Cont* 20°C/12°C (d/n) treatment for 3 days], *HT* 41°C/30°C (d/n) stress treatment for 3 days

**Fig.2.**
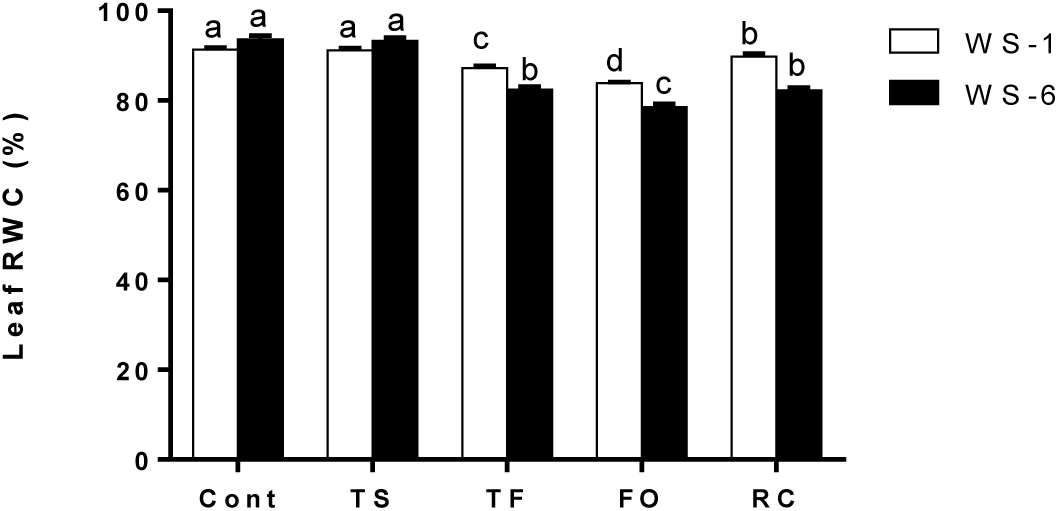
Effects of heat stress on leaf relative water content (RWC) in wucai plants. Values represent the mean ± SE (n=3). Letters indicate significant differences at P<0.05 according to Duncan’s multiple range tests. *Cont* 20°C/12°C (d/n) treatment for 3 days, *TS* 27°C/18°C (d/n) stress treatment for 3 days, *TF* 34°C/24°C(d/n) stress treatment for 3 days, *FO* 41°C/30°C (d/n) stress treatment for 3 days, *Recovery* afert *HT* treatment, transferred to 20°C/12°C (d/n) treatment for 3 days

### Chlorophyll content

In both wucai cultivars, the changes in Chl a, Chl b, Chl a+b and carotenoids contents showed the same trends in response to heat stress (Fig 3). Relative to the controls, Chl a, Chl b, Chl a+b and carotenoids contents was not significantly affected in WS-1 and WS-6 under 27°C, while at 34°C, they were deceased 3.26%, 4.00%, 3.52%, 6.03% and 6.18%, 5.91%, 6.10%, 11.44% respectively. As the temperature increased to 41°C, the Chl a, Chl b and Chl a+b contents were sharply decreased by 8.12%, 9.84%, 8.73%, 12.15% and 11. 57%, 15.08%, 12.78%, 23.60% in WS-1 and WS-6 respectively. Moreover, larger decreases in Chl a, Chl b, Chl a+b and carotenoids contents occurred at 34°C and 41°C in WS-6, compared with WS-1. After plants were transferred to moderate temperature condition for 3 days, average recovery rates in Chl a, Chl b, Chl a+b contents were 97.64%, 98.77%, 98.03%, 100.57% of WS-1 and below 90..06%, 88.04%, 89.36%, 91.89% of WS-6. While the recovery from 41°C heat stress, WS-1 occurred more quickly and mostly reached that of the control level.

**Fig.3.**
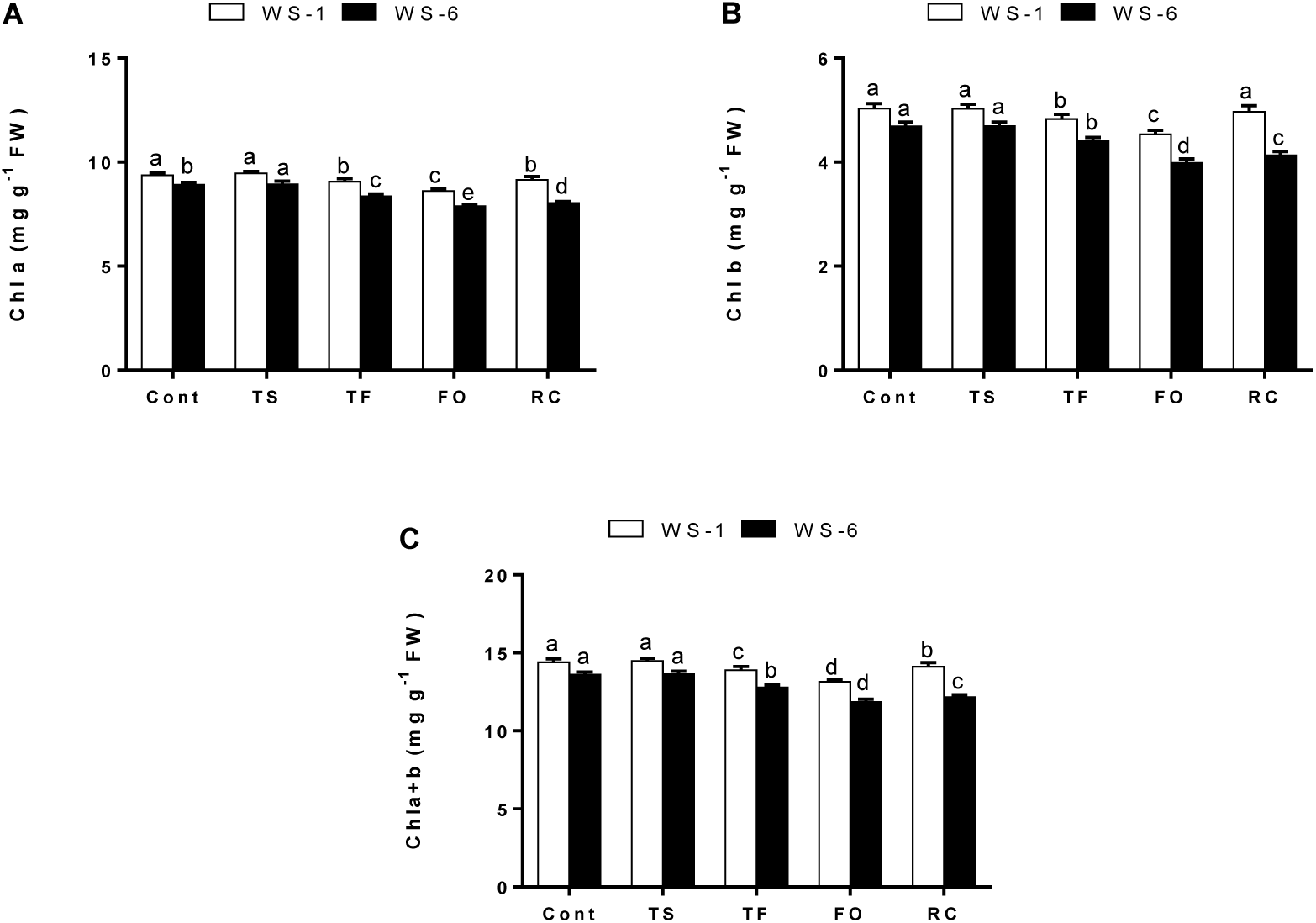
Effects of heat stress on Chl a, Chl b and Chl a+b contents in wucai plants. Values represent the mean ± SE (n=3). Letters indicate significant differences at P<0.05 according to Duncan’s multiple range tests. *Cont* 20°C/12°C (d/n) treatment for 3 days, *TS* 27°C/18°C (d/n) stress treatment for 3 days, *TF* 34°C/24°C(d/n) stress treatment for 3 days, *FO* 41°C/30°C (d/n) stress treatment for 3 days, *Recovery* afert *HT* treatment, transferred to 20°C/12°C (d/n) treatment for 3 days

### Photosynthetic parameters

P_N_ was unchanged in either cultivars at 27°C, but significantly decreased in both cultivars under 34°C and 41°C, relative to their respective controls (Fig 4 A). At 34°C, P_N_ was significantly decreased by 16.70% in WS-1, while decreased by 24.14% in WS-6. When the temperature increase to 41°C, larger decreases in PN occurred at 41°C decreased by 41.89% in WS-6 and decreased by 28.65% in WS-1, compared with their respectively controls. After recovery, average recovery rates were 92.00% of WS-1 and below 70.06% of WS-6.

**Fig.4.**
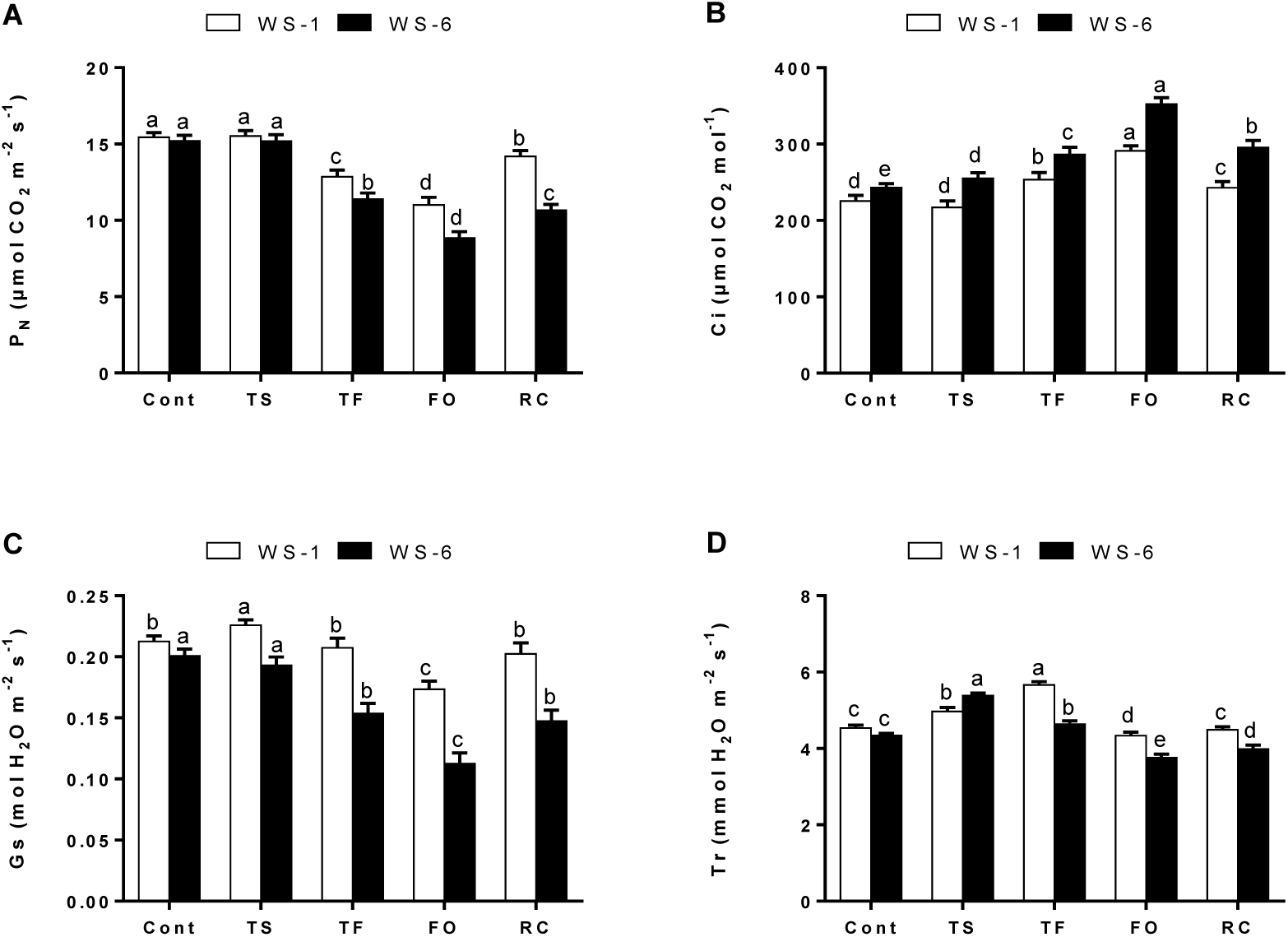
Effects of heat stress on net photosynthetic rate (P_N_), stomatal conductance (Gs), intercellular CO_2_ concentration (Ci), and transpiration rate (E) in wucai seedlings. Letters indicate significant differences at P<0.05 according to Duncan’s multiple range tests. *Cont* 20°C/12°C (d/n) treatment for 3 days, *TS* 27°C/18°C (d/n) stress treatment for 3 days, *TF* 34°C/24°C(d/n) stress treatment for 3 days, *FO* 41°C/30°C (d/n) stress treatment for 3 days, *Recovery* afert *HT* treatment, transferred to 20°C/12°C (d/n) treatment for 3 days

With the temperature increased, C_i_ of heat-sensitive WS-6 was increased significantly, while heat-tolerant WS-1 was unaffected by 34°C. At 41°C, C_i_ was significantly decreased by 29.12% in WS-1, while decreased by 44.90% in WS-6 (Fig 4 B). When the plants were transferred to moderate temperature condition for 3 days later, larger increases of C_i_ were presented in WS-6 of 21.46% relative to those in WS-1 of 7.58%.

The G_S_ values increased to maximum at 27°C and then declined with increasing temperature in WS-1, while decreased at 27°C in WS-6. Larger decreases of g_S_ were presented in WS-6 at 34°C and 41°C relative to those in WS-1 (Fig 4 C). Average recovery rates were 94.84% of WS-1 and below 73.13% of WS-6.

The response of *E* to heat stress were similar in both cultivars. They were significantly increased at 34°C and decreased at 41°C, relative to their respective controls (Fig 4 D). At 34°C, *E* was significantly increased by 24.84% in WS-1, while increased by 6.84% in WS-6. When the temperature increase to 41°C, larger decreases in *E* occurred at 41°C decreased by 13.43% in WS-6 and decreased by 4.36% in WS-1, compared with their respectively controls.

### Chlorophyll fluorescence parameters

A typical fast chlorophyll fluorescence rise kinetics shows a sequence of phases from the initial (F_0_) to the maximal (F_M_) fluorescence value, which have been labeled step O (20 μs), J (~2 ms), I (~30 ms), and P (equal to F_M_). At 20°C, both wucai cultivars showed a tipical O-J-I-P steps. Moreover, at 41°C, the K-step appeared in WS-6 cultivars, while WS-1 still showed a tipical O-J-I-P steps (Fig 5).

**Fig.5.**
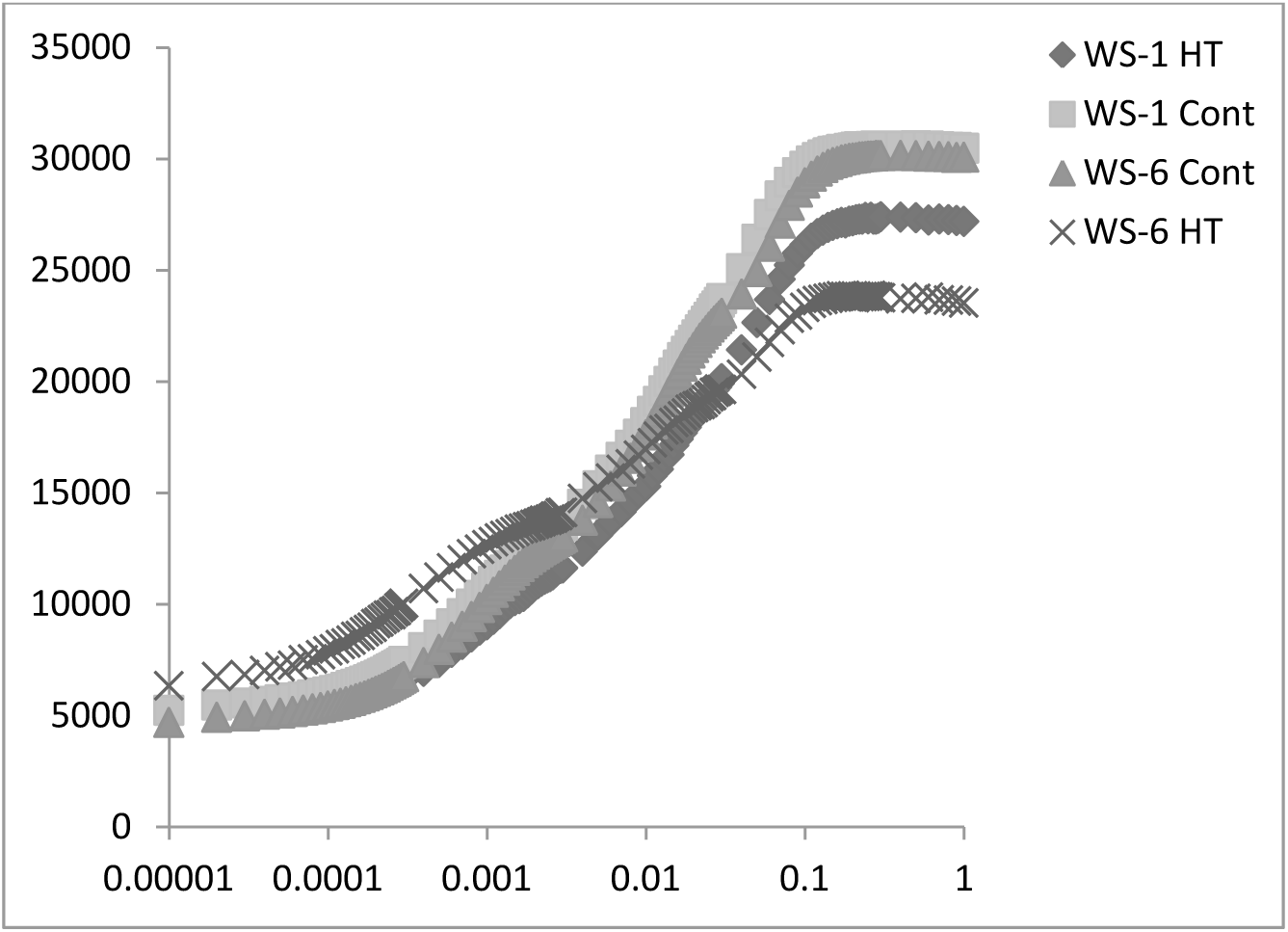
Effects of heat stress on O-J-I-P steps in wucai seedlings. *Cont* 20°C/12°C (d/n) treatment for 3 days, *HT* 41°C/30°C (d/n) stress treatment for 3 days

The F_0_ values of both cultivars were significantly increased at higher temperatures, especially in WS-6. At 41°C, F_0_ values were, respectively, increased by 23.94% and 34.33% in WS-1 and WS-6, relative to their controls.

WS-1 had similar trend of F_V_/F_M_ that of WS-6 under normal condition in leaves (Fig 6 A). At 27°C, F_v_/F_m_ values were both increased, but had no significantly differences to their controls. However, 34°C and 41°C heat stress resulted in declines in F_V_/F_M_ of two cultivars (Fig 6 B). Compared to their respective controls, the F_V_/F_M_ in WS-1 were decreased by 5.75% and 9.91%; those of WS-6 were decreased by 9.39% and 27.30%. The decrease extent of F_V_/F_M_ of WS-6 was higher than that of WS-1.

**Fig.6.**
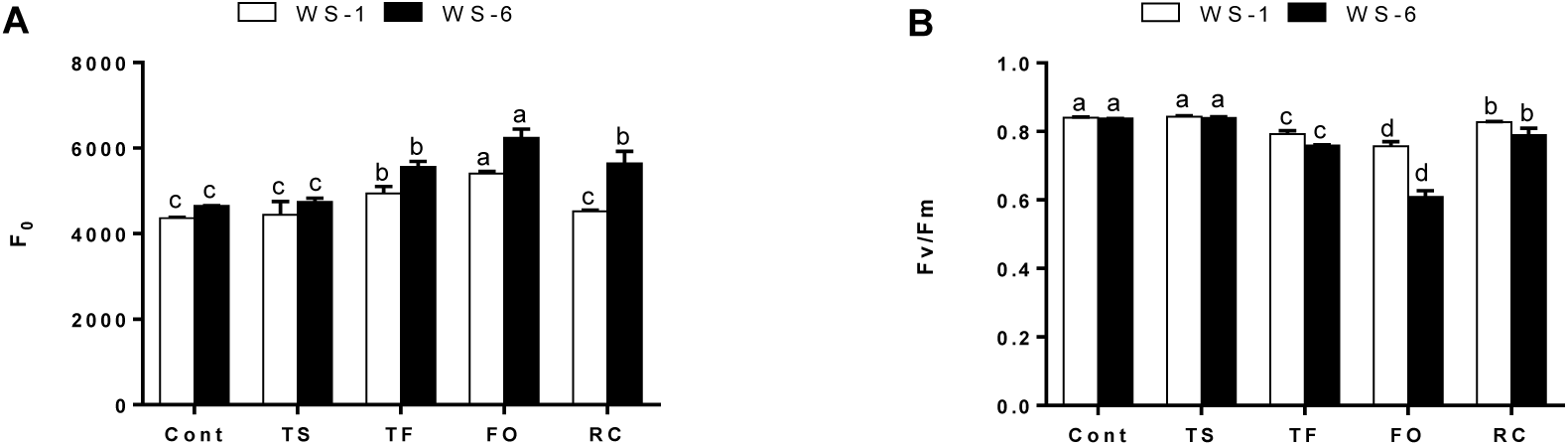
Effects of heat stress on F_0_ and F_V_/F_M_ in wucai seedlings. Letters indicate significant differences at P<0.05 according to Duncan’s multiple range tests. *Cont* 20°C/12°C (d/n) treatment for 3 days, *TS* 27°C/18°C (d/n) stress treatment for 3 days, *TF* 34°C/24°C(d/n) stress treatment for 3 days, *FO* 41°C/30°C (d/n) stress treatment for 3 days, *Recovery* afert *HT* treatment, transferred to 20°C/12°C (d/n) treatment for 3 days

Moreover, the F_0_ values and F_V_/F_M_ values in WS-1 was similar to that of its control after recovery, while WS-6 had obviously increases of 21.43% and 5.77% respectively (Fig 6). Based on RC, there were no significanly changes in ABS/RC, TR_0_/RC, DI_0_/RC, ET_0_/RC for either cultivars at 27°C(Fig 7). At high temperature, ABS/RC, TR_0_/RC, DI_0_/RC were significantly increased, whlie ET_0_/RC were decreased. At 41°C, ABS/RC of WS-1 and in WS-6 were, respectively, increased by 26.12% and 48.13%, TR_0_/RC of WS-1 and in WS-6 were, respectively, increased by 31.54% and 61.03%, DI_0_/RC of WS-1 and in WS-6 were, respectively, increased by 64.41% and 122.42%, and ET_0_/RC of WS-1 and in WS-6 were, respectively, decreased by 15.11% and 28.62%.

**Fig.7.**
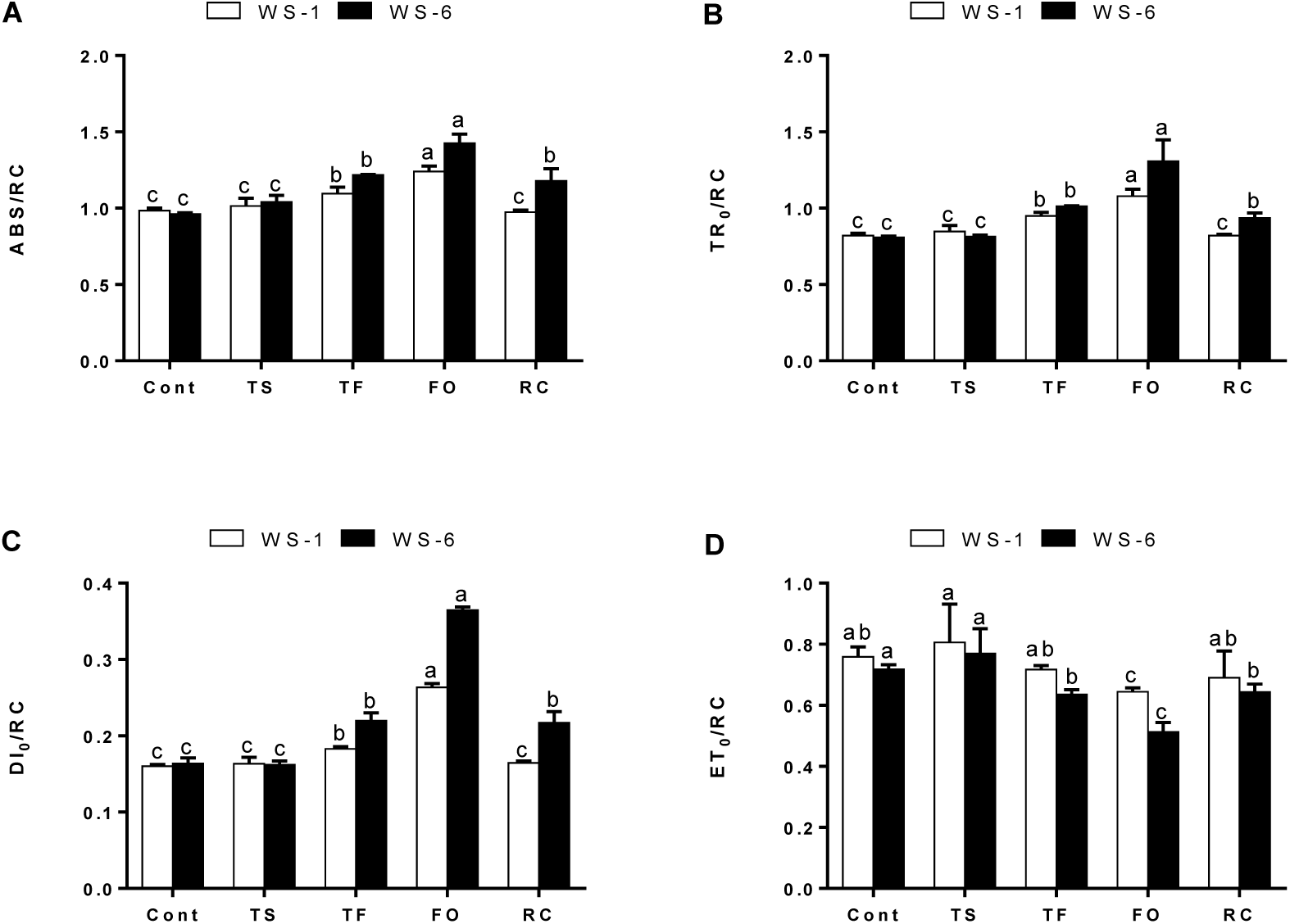
Effects of heat stress on ABS/RC, TR_0_/RC, DI_0_/RC and ET_0_/RC in wucai seedlings. Letters indicate significant differences at P<0.05 according to Duncan’s multiple range tests. *Cont* 20°C/12°C (d/n) treatment for 3 days, *TS* 27°C/18°C (d/n) stress treatment for 3 days, *TF* 34°C/24°C(d/n) stress treatment for 3 days, *FO* 41°C/30°C (d/n) stress treatment for 3 days, *Recovery* afert *HT* treatment, transferred to 20°C/12°C (d/n) treatment for 3 days

Once stress conditions were removed, ABS/RC, TR_0_/RC, DI_0_/RC, ET_0/_RC of WS-1 were almost recovered to the control level, while of WS-6 were still had significantly differences compared with their controls (Fig 7).

The RC/ CS_M_ values of both cultivars were significantly decreased at higher temperatures, especially in WS-6 (Fig 8). There were no significanly changes for either cultivars at 27°C. But at 34°C, RC/ CS_M_ values were, respectively, decreased by 10.63% and 15.42% in WS-1 and WS-6, relative to their controls. With the temperature increased to 41°C, RC/ CS_M_ values were, respectively, decreased by 10.63% and 15.42% in WS-1 and WS-6, relative to their controls. After recovery, the values of WS-1 were reached the control level, while the values of WS-6 still have large difference compared to the control.

**Fig.8.**
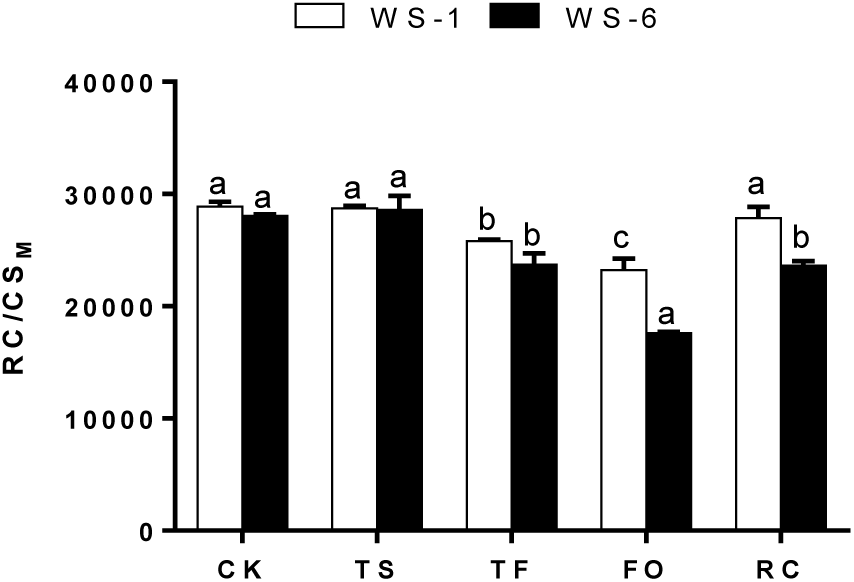
Effects of heat stress on RC/ CS_M_ in wucai seedlings. Letters indicate significant differences at P<0.05 according to Duncan’s multiple range tests. *Cont* 20°C/12°C (d/n) treatment for 3 days, *TS* 27°C/18°C (d/n) stress treatment for 3 days, *TF* 34°C/24°C(d/n) stress treatment for 3 days, *FO* 41°C/30°C (d/n) stress treatment for 3 days, *Recovery* afert *HT* treatment, transferred to 20°C/12°C (d/n) treatment for 3 days

Based on CS_M_, there were no significanly changes in ABS/CS_M_, DI_0_/CS_M_, ET_0_/ CS_M_ for either cultivars at 27°C. But the TR_0_/CS_M_ were decreased at 27°C in both wucai cultivars. At high temperature, ABS/CS_M_, TR_0_/CS_M_, DI_0_/ CS_M_ were significantly increased, whlie ET_0_/ CS_M_ were decreased (Fig 9). At 41°C, ABS/ CS_M_, TR_0_/CS_M_, ET_0_/ CS_M_ of WS-1 were, respectively, decreased by 14.75%, 24.29% and 21.78% in WS-1, while were respectively decreased by 28.48%, 41.34%, 50.61% in WS-6. Additionally, DI_0_/CS_M_ of WS-1 and in WS-6 were, respectively, increased by 18.49% and 34.33% under 41°C heat stress.

**Fig.9.**
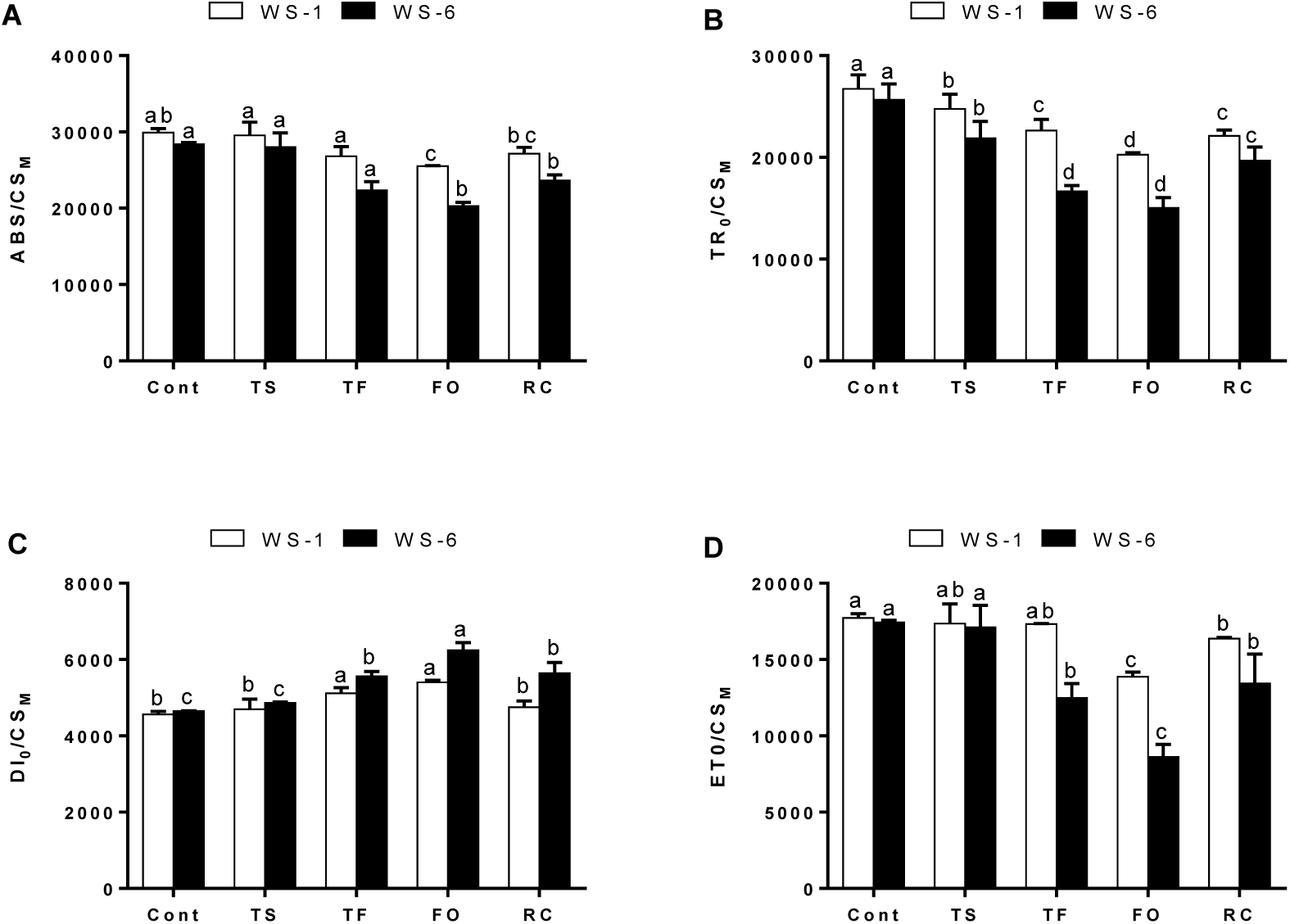
Effects of heat stress on ABS/ CS_M_, TR_0_/ CS_M_, DI_0_/ CS_M_, ET_O_/ CS_M_ in wucai seedlings. Letters indicate significant differences at P<0.05 according to Duncan’s multiple range tests. *Cont* 20°C/12°C (d/n) treatment for 3 days, *TS* 27°C/18°C (d/n) stress treatment for 3 days, *TF* 34°C/24°C(d/n) stress treatment for 3 days, *FO* 41°C/30°C (d/n) stress treatment for 3 days, *Recovery* afert *HT* treatment, transferred to 20°C/12°C (d/n) treatment for 3 days

The levels of ABS/ CS_M_ and DI_0_/ CS_M_ in WS-1 returned to the control values when plants subjected to heat stress were transferred to the suitable environment for 3 days. However, the levels of TR_0_/CS_M_, ET_0_/CS_M_ in WS-1 and ABS/ CS_M_, TR_0_/ CS_M_, DI_0_/CS_M_, ET_0_/CS_M_ in WS-6 were still had significantly differences compared with their controls (Fig 9).

PI_ABS_ is a very sensitive indicator of the physiological status of plants subjected to environmental stress. Following heat stress, PI_ABS_ was decreased in both varieties. The PI_ABS_ values of both cultivars were sharply decreased at 41°C, especially in WS-6, decreased by 71.50% compared to the control (Fig 10). Moreover, after translated to the control level, WS-1 had higher recovery rate than WS-6.

**Fig.10.**
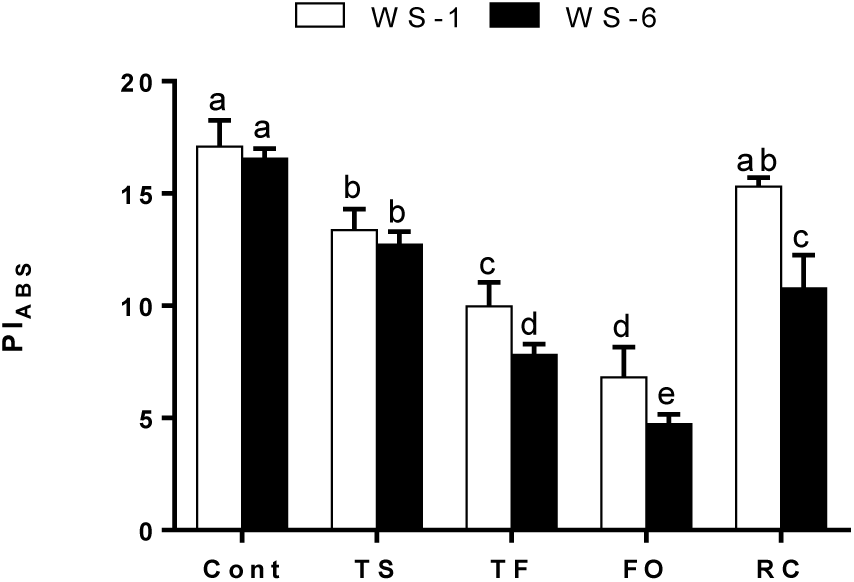
Effects of heat stress on PI_ABS_ in wucai seedlings. Letters indicate significant differences at P<0.05 according to Duncan’s multiple range tests. *Cont* 20°C/12°C (d/n) treatment for 3 days, *TS* 27°C/18°C (d/n) stress treatment for 3 days, *TF* 34°C/24°C(d/n) stress treatment for 3 days, *FO* 41°C/30°C (d/n) stress treatment for 3 days, *Recovery* afert *HT* treatment, transferred to 20°C/12°C (d/n) treatment for 3 days

### Fatty acid composition of thylakoid membranes

The fatty acids of thylakoid membrane lipids of wucai leaves were separated and analyzed by gas chromatography. As shown in Table 2, two saturated fatty acids (palmitic acid, and stearic acid) and four unsaturated ones (palmitoleic acid, oleic acid, linoleic acid, and linolenic acid) were the main fatty acids among the thylakoid membrane lipids. Specifically, in response to high temperature treatment, in WS-1, palmitic and stearic acid content was increased by 55.35% and 65.20%, whereas palmitoleic acid, oleic acid, linoleic acid, and linolenic acid content was decreased by 11.47%, 30.01%, 18.63% and 34.66% respectively, compared with control plants. While in WS-6, palmitic and stearic acid content was increased by 88.80% and 31.96%, whereas palmitoleic acid, oleic acid, linoleic acid, and linolenic acid content was decreased by 1.41%, 4.80%, 3.32% and 22.50% respectively, compared with control plants. When exposed to heat stress for 3 days, High temperature stress significantly increased saturated fatty acid levels, but decreased all of the unsaturated fatty acid content, the ratio of unsaturated to saturated fatty acids and IUFA. After plants were transferred to moderate temperature condition for 3 days, the ratio of unsaturated to saturated fatty acids and IUFA respectively average recovery rates were 90.38% and 96.45% of WS-1 plants and below 76.32% and 88.89% of WS-6 plants. All the results suggest that the plant alleviates heat stress-induced thylakoid membrane lipid peroxidation by enhancing unsaturated fatty acid content.

**Table 2.** Effects of heat stress on the fatty acid composition of thylakoid membranes in two wucai cultivars

| Fatty acid Composition<br>(mol %) | Treatments |  |  |  |  |  |
| --- | --- | --- | --- | --- | --- | --- |
|  | WS-1 Cont | WS-1 HT | WS-1 Recovery | WS-6 Cont | WS-6 HT | WS-6 Recovery |
| Palmitic acid(16:0) | 23.042±0.469b | 35.795±0.715a | 24.154±0.715b | 23.873±0.390c | 29.315±0.774a | 27.653±0.708b |
| Stearic acid(18:0) | 9.221±0.316c | 15.235±0.763a | 10.362±0.434b | 10.353±0.362c | 13.662±0.639a | 12.878±0.781b |
| Palmitoleic acid(16:1) | 10.124±0.364a | 8.963±0.513c | 9.584±0.614b | 9.910±0.548a | 9.770±0.770a | 9.224±0.595b |
| Oleic acid(18:1) | 12.381±0.443a | 8.665±0.399b | 12.450±0.681a | 11.211±0.381a | 10.673±0.579b | 10.263±0.672b |
| Linoleic acid(18:2) | 11.222±0.349a | 9.131±0.595b | 10.981±0.616a | 10.432±0.330a | 10.086±0.469b | 10.211±0.501ab |
| Linolenic acid(18:3) | 34.021±0.671a | 22.231±0.723c | 32.482±0.640b | 34.222±0.531a | 26.522±0.652c | 29.781±0.663b |
| UFA | 67.748±1.729a | 48.990±1.980c | 65.497±2.150b | 65.785±1.585a | 57.051±2.081c | 59.479±2.072b |
| SFA | 32.264±0.508c | 51.030±1.409a | 34.516±1.105b | 34.226±0.583c | 42.977±1.096a | 40.532±1.358b |
| UFA/SFA | 2.100±0.062a | 0.960±0.012c | 1.898±0.026b | 1.922±0.140a | 1.327±0.019c | 1.467±0.004b |
| IUFA | 147.012±3.263a | 102.582±3.882c | 141.443±4.169b | 144.661±2.806a | 120.181±3.671c | 129.251±3.626b |

### Blue native (BN) /SDS-PAGE electrophoresis

To identify heat stress mediated thylakoid membrane proteins, we carried out a comparative proteomic analysis of thylakoid membranes after 3 days of treatment. Protein complexes were first solubilized from thylakoid membranes and then separated by BN-PAGE. After the first dimensional separation, seven major protein complexes were obtained (Fig 11). Mass spectrometric analysis identified the complexes as supercomplexes, PSI-LHCII/PSII monomer, PSII monomer, CP43 less PSII/ATP synthase, LHCII trimer, LHCII monomer and ATP synthase. With temperature increased, heat stress decreased levels of PSII protein complex, monomeric LHCII bands and ATP synthase bands, but increased trimeric LHCII bands. WS-1 had slightly change compared to WS-6 under high temperature stress (Fig 12).

**Fig.11.**
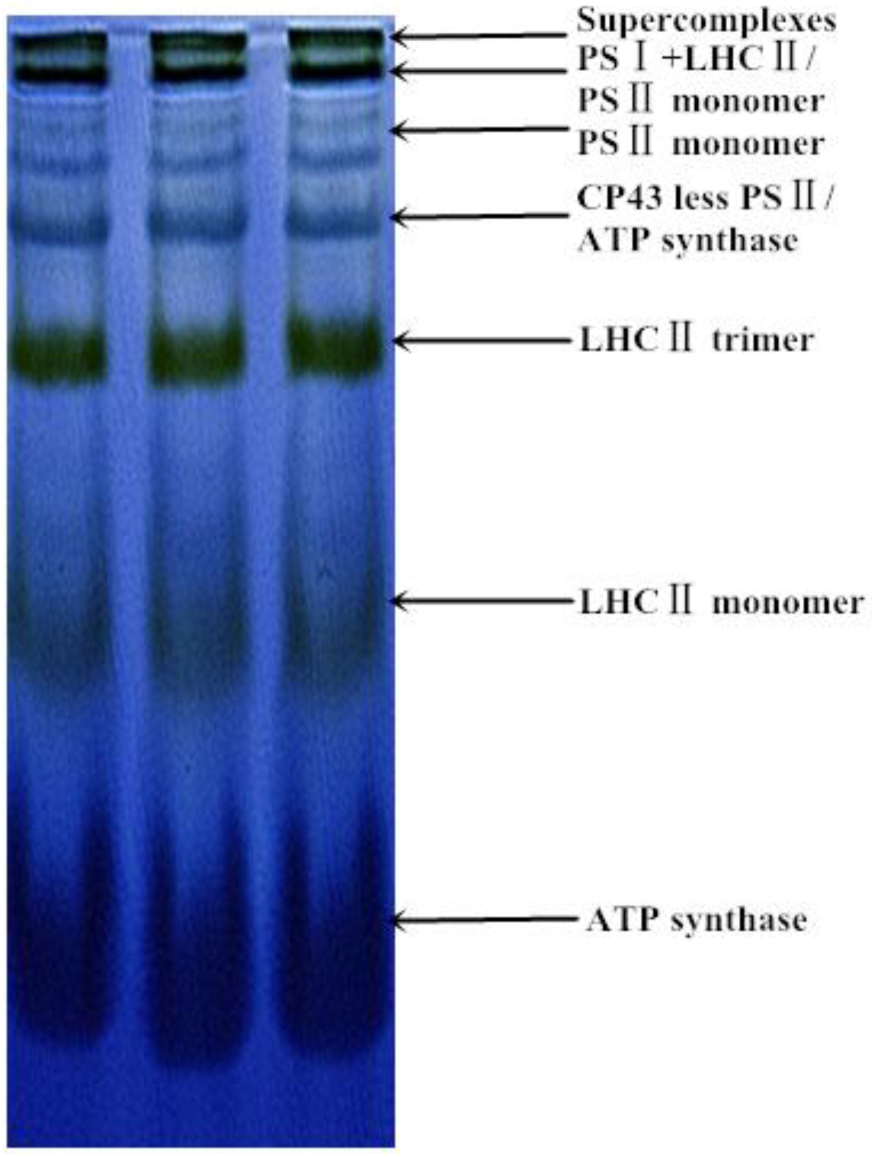
Membrane protein complexes in wucai seedlings. Blue-native gel electrophoresis of membrane complexes protein from stroma thylakoids were solubilised by n-dodecylmaltoside. Protein complexes were first solubilized from thylakoid membranes using n-dodecyl-ß-D-maltoside (DM) and then separated by BN-PAGE. After the first dimensional separation, seven major protein complexes were obtained.

**Fig.12.**
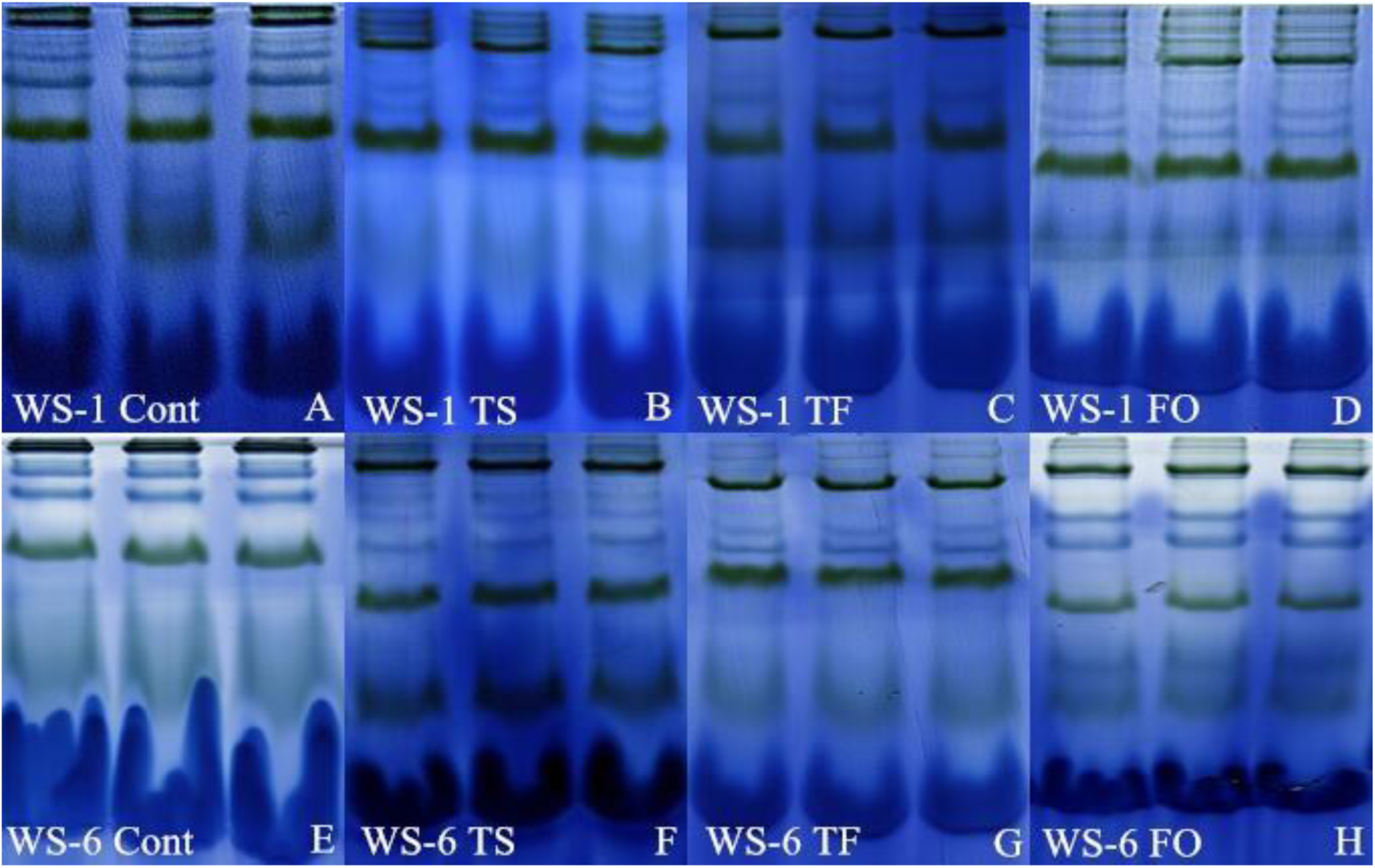
Effects of heat stress on membrane protein complexes in wucai seedlings. *Cont* 20°C/12°C (d/n) treatment for 3 days, *TS* 27°C/18°C (d/n) stress treatment for 3 days, *TF* 34°C/24°C(d/n) stress treatment for 3 days, *FO* 41°C/30°C (d/n) stress treatment for 3 days

To fingerprint these complexes, BN-PAGE gels were excised and layered onto PAGE gel slabs and then subjected to SDS-urea-PAGE followed by Coomassie blue G-250 staining. This separation enabled visualization of the subunit patterns of the complexes. Analysis of the BN-PAGE gel using ImageMaster 2D Platinum software revealed more than 60 CBB-stained protein spots associated with molecular masses of 14.4–116 kDa. Of these, 15 protein spots differentially regulated in response to WS-1 and WS-6 under heat stress were excised from the gels and identified (Fig 13). Differential proteins including PS I P700 chlorophyll, chlorphyll a-b binding, PS II, D2, CP43, PS I, ATP synthase beta subunit and ATP synthase alpha subunit (Table 4).

**Fig.13.**
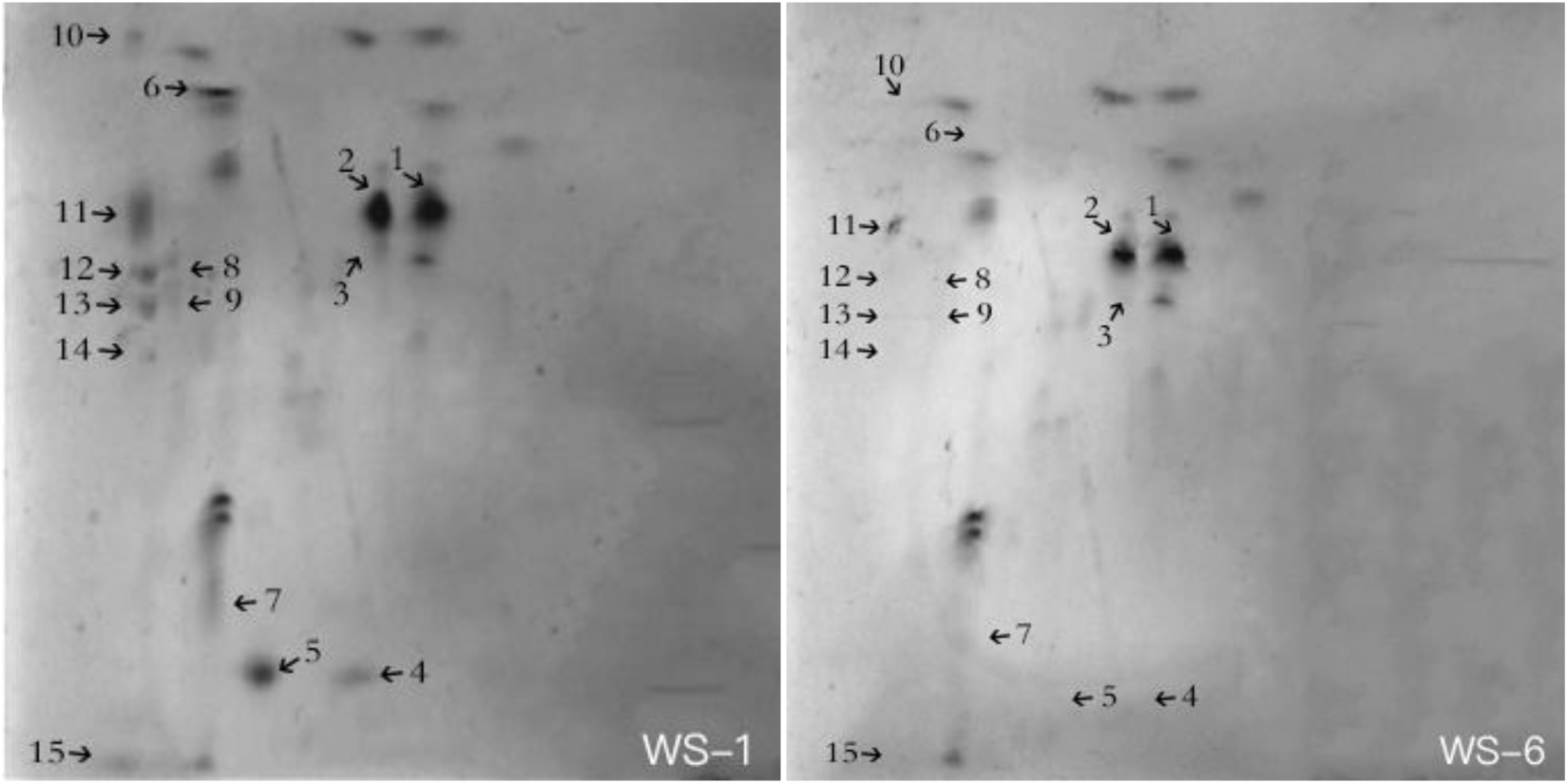
Effects of heat stress on differential membrane proteins in wucai seedlings. 41°C/30°C (d/n) stress treatment for 3 days

**Table 3.** Derivations and definitions of JIP parameters directly obtained from the recorded OJIP fluorescence transients

| Technical fluorescence parameters | Definition |
| --- | --- |
| $F_0$ | Fluorescence value at 50 ms, used as initial value of fluorescence |
| $F_V/F_M$ | Maximum quantum yield of primary PSII photochemistry |
| $ABS/RC$ | Average absorbed photon flux per PSII reaction centre |
| $TR_O/RC$ | Maximum trapped exciton flux per PSII reaction centre |
| $DI_O/RC$ | Dissipated energy flux per PSII reaction centre |
| $ET_O/RC$ | Electron transport flux from $Q_A$ to $Q_B$ per PSII reaction centre |
| $RC/CS_M$ | Number of active reaction centres per cross section of PSII |
| $ABS/CS_M$ | Average absorbed photon flux per cross section of PSII |
| $TR_O/CS_M$ | Maximum trapped exciton flux per cross section of PSII |
| $DI_O/CS_M$ | Dissipated energy flux per cross section of PSII |
| $ET_O/CS_M$ | Electron transport flux from $Q_A$ to $Q_B$ per cross section of PSII |
| $PI_{ABS}$ | Performance index on absorption basis |

**Table 4.** Differentially synthesised proteins identified by MALDI-TOF-MS and LC-ESI-MS/MS from the spots in BN/SDS-PAGE in Fig. 13.

| Spots | Protein name | Source | Accession No |
| --- | --- | --- | --- |
| 1 | Photosystem II CP43 reaction center protein OS | Brassica oleracea var. capitata | tr\|A0A0H3XZM8\|A0A0H3XZM8_B<br>RAOC |
| 2 | ATP synthase subunit alpha OS | Brassica napus | tr\|A0A078G6X6\|A0A078G6X6_BR<br>ANA |
| 3 | BnaC03g29960D protein OS | Brassica napus | tr\|A0A078DQX4\|A0A078DQX4_BR<br>ANA |
| 4 | Chlorophyll a-b binding protein, chloroplastic (Fragment) OS | Sinapis alba | tr\|A0A075C5B7\|A0A075C5B7_SIN<br>AL |
| 5 | Photosystem II D2 protein OS | Brassica oleracea var. oleracea | tr\|A0A0D3BE55\|A0A0D3BE55_BR<br>AOL |
| 6 | Photosystem II OS | Brassica oleracea var. capitata | tr\|A0A0H3XZM8\|A0A0H3XZM8_B<br>RAOC |
| 7 | Chlorophyll a-b binding protein, chloroplastic (Fragment) OS | Sinapis alba | tr\|A0A075C5B7\|A0A075C5B7_SIN<br>AL |
| 8 | Photosystem I P700 chlorophyll a apoprotein A2 OS | Brassica oleracea var. capitata | tr\|A0A0H3Y227\|A0A0H3Y227_BR<br>AOC |
| 9 | Chlorophyll a-b binding protein, chloroplastic (Fragment) | Brassica napus | tr\|A0A075C5B7\|A0A075C5B7_SIN<br>AL |
| 10 | Chlorophyll a-b binding protein, chloroplastic OS | Brassica oleracea var. oleracea | tr\|A0A0D3AHD4\|A0A0D3AHD4_B<br>RAOL |
| 11 | ATP synthase subunit beta, chloroplastic OS | Raphanus sativus var. raphanistroides | tr\|A0A249RSF6\|A0A249RSF6_RAP<br>SA |
| 12 | Chlorophyll a-b binding protein, chloroplastic OS | Brassica napus | tr\|A0A078J8I6\|A0A078J8I6_BRAN<br>A |
| 13 | Chlorophyll a-b binding protein, chloroplastic (Fragment) OS | Sinapis alba | tr\|A0A075C5B7\|A0A075C5B7_SIN<br>AL |
| 14 | Chlorophyll a-b binding protein, chloroplastic OS | Brassica napus | tr\|A0A078FKQ7\|A0A078FKQ7_BR<br>ANA |
| 15 | Chlorophyll a-b binding protein, chloroplastic OS | Brassica oleracea var. oleracea | tr\|A0A0D3AHD4\|A0A0D3AHD4_B<br>RAOL |

## Discussion

It is well known that high temperature can severly inhibited plant growth. It was reported that heat stress resulted in reduction on plant height, stem diameter, leaf width, leaf length and biomass in plants (Garruna-Hernández et al., 2014; Nayyar et al., 2014; Siddiqui et al., 2015). In this study, wucai cultivars responded comprehensively in terms of growth and physiological characteristics and differential resistance to heat stress. The heat stress exhibited a negative effect on both wucai cultivars. Plant height, stem diameter and single plant weight (fresh weight and dry weight) were significantly decreased under high temperature treatment (Table 1). This might be caused by a reduction in the relative growth rates of stalk and stem due to decreased net assilimilation and the loss of turgidity and RWC (Fig 2) under heat stress(Wahid 2007., Srivastava et al. 2012). Leaf RWC has been established as an indicator of state of water balance of plants essentially in terms of the physiological consequences of cellular water deficit (Wahid and Close 2007, Kumar et al. 2008, Kesici et al., 2013). In both cultivars, RWC declined linearly from ambient to modest temperature. Among the cultivars, WS-1 exhibited the higher RWC while WS-6 had the lower RWC under heat stress (Fig 2). The high level of temperature stress might be altered cell division and cell elongation resulted in reduced leaf length and leaf width in both cultivars (Table 1). In this study, larger morphological changes occurred in WS-6 than in WS-1, indicating that the effects of high temperatures on fresh weight, dry weight, plant height, stem diameter, leaf width and length were cultivar dependent in wucai (Table 1).

In several studies, heat stress led to reductions in photosynthetic pigment contents (Marchand et al. 2005) and impaired Chl biosynthesis in plastids (Dutta et al. 2009). In both cultivars, Total Chl synthesis was impaired with increasing temperature (Fig 3). The decrease in total Chl biosynthesis in wucai cultivars may be due to increased activity of the chloroplast degrading enzyme, induced changes in ultrastructure of chloroplasts, and the chloroplasts could gradually lose its capacity to capture radiation energy (Kudoh and Sonoike 2002). Also, heat stress could cause lipid peroxidation of chloroplasts and thylakoid membranes (Mohammed and Tarpley 2010) and various changes in chloroplasts (Lipiec et al., 2013). Moreover, WS-6 had a higher decline in chlorophyll content than WS-1 under each of high temperature treatments (Fig 3). Because of total Chl is an important biomolecule in plants photosynthesis. Additional, in our stduy, the different level of photosynthetic parameters and chlorophyll fluorescence parameters may due to different Chl content in two wucai cultivars under heat stress (Fig 4–10). These results suggest that the heat-tolerant cultivar has better self-protection processes for resisting heat stress.

Several studies have shown that photosynthesis is very sensitive to high temperatures, and is often the first cellular function to be impaired by heat stress (Tan et al. 2011; Greer and Weedon 2012). Furthermore, plants grown under temperature stress usually show a decrease in photosynthesis due to stomatal closure, chloroplast impairment, or limitation of the carbon assimilation (Pastenes and Horton 1996, Wise and Olson 2004). In this study, the high temperature stress, especially 41°C, caused decreases in P_N_, G_S_, and E, whereas the C_i_ value was increased after 3 days (Fig 4). These results indicated that the high temperature caused the reduction of PN and this reduction was mainly caused by non-stomatal limitation under severe heat stress due to higher C_i_ with lower G_s_ (Ploschuk et al. 2014), but stomatal limitation would occurred under milder stress conditions (Wu et al. 2001). Moreover, photosynthetic parameters exhibited greater changes in WS-6 than in WS-1 under heat stress. Our results confirmed that the heat stress mediated decline of PN was also cultivar dependent. Compared with the heat-sensitive cultivar, the heat-tolerant cultivar was better able to regulate photosynthesis under heat stress. Net photosynthetic rate in WS-6 had lower recovery ratios compared with WS-1 indicating that WS-1 was more sensitive to the temperature stress than WS-6.

One of the most sensitive components of the photosynthetic apparatus in plants to high temperatures is PSII (Srivastava et al. 1997), and it is considered to play a key role in photosynthesis under environmental stress (Baker and Rosenqvist 2004). PSII activity was greatly reduced or completely arrested under heat stress (Morales et al. 2003). Heat stress could caused F_0_ increses duo to the physical separation of the PSII RCs from their associated pigment antennae resulting in blocked energy transfer to the PSII traps (Fig 6 A). Thus, heat inactivation of PSII may be accompanied by the aggregation and subsequent dissociation of the light-harvesting complex (Li et al., 2009). In our study, larger decreases in Fv/Fm were at higher temperatures (Fig 6 B), implying that photoinhibition occurred under heat stress and was magnified by higher temperatures. The degree of photoinhibition appeared to be greater in the heat-sensitive cultivar than in the heat-tolerant one.

A typical fast chlorophyll fluorescence rise kinetics shows a sequence of phases from the initial (F_O_) to the maximal (F_M_) fluorescence value, which have been labeled step O (20 μs, all RCs open), J (~2 ms), I (~30 ms), and P (equal to F_M_ when all RCs are closed) (Strasser and Strasser 1995). Besides the basic O-J-I-P steps, others also appear in certain conditions, such as the K-step, relating to the inactivation of the oxygen-evolving complex (OEC)(Tsimilli-Michael et al., 1999; Strasser et al., 2004). On the other hand, one to two of the basic O-J-I-P steps will disappear in some stress situations. Under strong heat stress (above 44°C), the J- and I-steps disappear with a concomitant appearance of the K peak as a predominant step in fluorescence rise kinetics because the OEC has been damaged completely (Strasser et al., 2004; Chen et al., 2016). Our experiments show that appearance of the K-step is the major change in fast chlorophyll fluorescence rise kinetics of croftonweed leaves exposed to high temperature (Fig 5). The phenomenological appearance of the K-step is a typical characteristic of the fluorescence rise kinetics in heated-samples, which is specifically attributed to the OEC destruction by release of the manganese cluster (Strasser et al., 2004). A high temperature can cause inactivation of OEC, inhibition of electron transport, and decrease in PSII photochemical efficiency (De Ronde et al., 2004; Wahid et al., 2007; Sharkey et al. 2005, Allakhverdiev et al. 2008). It is reported that heat stress leads to the dissociation of the OEC causing an imbalance between the electron flow from the OEC to the RC and towards PSII acceptor side, the alternative internal electron donor such as proline can donate electrons to PSII instead of H_2_O (De Ronde et al. 2004; Oukarroum et al., 2013). This will result in a short-lived increase in the reduced Pheo/Q_A_ concentration, creating a K-peak appearing at about 300 μs. Hence, the increasing amplitude of the K-step or ∆K peak is associated with the OEC injury degree (Brestic ea al. 2013; Strasser, 1997; Oukarroum et al. 2016). In fact, the OEC with manganese cluster is very sensitive to heat stress, and the OEC damage is one of the earliest events affected by heat stress. While the appearance of a conspicuous K peak requires higher temperature intensities (above 40°C) or long heat duration in moderate temperature at 40°C. Just above 40°C high temperatures, a significant increase of the level of K-step (V_K_ or W_K_) or ∆K peak (∆W_K_) was observed(Chen et al., 2016). In our study, tolerant WS-1 plants had a less increase of the ∆K peak than sensitive WS-6 plants at different heat stress level (Fig 5).

In addition to the widely used F_0_ and Fv/F_M_ parameter, we studied the JIP-test parameters ABS/CS_M_, DI_0_/CS_M_, TR_0_/CS_M_, RC/CS_M_ and ET_0_/CS_M_ which can be used to explain the stepwise flow of energy through PSII at the crosssection for maximum fluorescence (CS_M_) level. According to Sinsawat et al. (2004), decreased F_V_/F_M_ was primarily due to the decrease in ABS/CS, ET_0_/CS, TR_0_/CS and the increase in DI_0_/CS. A significant decrease in RC/CS, ABS/CS and TR_0_/CS further demonstrates that high temperatures indeed inactivated PSII RCs, reduced the function antenna size, and declined the specific rate of the exciton trapped by open RCs (Fig 8). We also notice that partial inactivation of PSII RCs in WS-6 plants already starts in mild heat stress at 34°C, moreover, complete inactivation of PSII RCs happens at 41°C severe elevated temperature (Fig 8). In this study, ABS/CS, ET_0_/CS, TR_0_/CS significantly decreased under heat stress, while DI_0_/CS increased (Fig 9), which was in accordance with previous studies (Song et al. 2013; Luo et al. 2014). It is proposed that the decrease of TR_0_/CS is mainly attributed to heat inactivation of RCs due to the dissociation of the manganese-stabilizing extrinsic 33 kDa protein from the PSII reaction center complex (Enami et al., 1994). Moreover, the increase in DI_0_/CS was larger in WS-1 than in WS-6. DI_0_/CS has been shown to be closely associated with the onset of harmless dissipation of excess energy (Gilmore 1997). The structural and functional aspects of PSII are interrelated. Under heat stress, the damage to the photosynthetic machinery will greatly affect the energetic cooperativity between the PSII units. In addition to the JIP-test parameters based on CS, we studied the others JIP-test parameters based on RC. In our study, heat stress reduced ET_0_/RC and increased DI_0_/RC, ABS/RC and TR_0_/RC detected in both cultivars(Fig 7). A consistent increase in inhibition of Q_A_^−^ re-oxidation (TR_0_/RC) and apparent antenna size, due to inactivation of some active RCs (ABS/RC) associated with accumulation of inactive RCs and an increase of DI_0_/RC was shown. DI_0_/RC has been previously related to high NPQ (Strasser et al. 2000; Ajigboye., et al.2016).

Our data showing a temperature-dependent linear decrease in the performance index PIABS of WS-6 plants suggested that heat treatment results in a significant decrease of the overall photosynthetic activity (Fig. 10). In contrast, a higher vitality is maintained in tolerant WS-1 plants heated by increased high temperatures (Fig 10). PI_ABS_ indeed can be regarded as a standard to successfully identify heat sensitivity of different croftonweed populations since it is the most sensitive experimentally derived parameter to various stress conditions. And the dramatic lowering of the overall photosynthetic activity of PSII (PIABS) should be attributed to inactivation of RCs (RC/CS) and inhibition of light reactions(Chen et al.2016, Strasser et al. 2004).

As the primary architecture of the cell, the membrane plays important roles in sensing environmental change, signal transduction and substance metabolism (Mittler et al., 2012; Guyot et al., 2015). High temperature increase the fluidity of the cytoplasmic membrane. To maintain membrane fluidity within an optimal range for normal biological activity, fatty acid desaturase genes which convert unsaturated fatty acids into saturated fatty acids, are down-regulated to decrease lipid unsaturation and thus decrease membrane fluidity in response to temperature up-shifts (de Mendoza, 2014; Holthuis and Menon, 2014). As cell membrane components, unsaturated fatty acids play key roles in the fluidity of cellular membranes (Mansilla et al., 2008; Ma et al., 2015). Alter the ratio of unsaturated to saturated fatty acids was responsible for the maintenance of membrane integrity and fluidity (Shu et al. 2015; Liu et al. 2017). In this study, the reduction of unsaturated fatty acid content of thylakoid membranes in both cultivars increased membrane liquidity and maintained an orderly arrangement of thylakoids. The total amount of UFA decreased while the total amount of SFA increased, yielding a lower UFA/SFA ratio that is indicative of increased membrane fluidity (Table 2). Our study confirmed that changes in unsaturated fatty acids composition can improve plant tolerance against heat stress.

The deleterious effects of heat stress on thylakoid membrane proteins is a mechanism protecting the photosynthetic apparatus from heat-induced damage. We performed BN/SDS-PAGE electrophoresis of thylakoid membrane fractions to detect major heat-induced modifications of protein structure (Fig 11). Membrane proteins responsible for maintained ability to produce energy and metabolism, the maintenance of efficient photorespiratory pathways, and antioxidant metabolism could serve important roles in regulating leaf senescence and whole-plant responses to heat stress in plants (Busby,. et al. 2006). Previous study reported that stress may interact not only with the PSII core proteins D1, D2, and CP43, but also with the LHCII antenna complex, PSI reaction centre proteins, and the ATP synthase subunit (Alfonso,. et al. 2001; Shu,. et al. 2015). Several other studies confirms that the transcription and translation of the LHCII supercomplex and the ATP synthase complex affected by regulating the energy balance of thylakoid membranes and in ensuring sophisticated coordination of energy excitation and dissipation (Hamdani,. et al. 2011; Hubbart,. et al. 2012). The protective action on PSII proteins can be explained in the modulation of synthesis and turnover of these proteins (Xue,. et al. 2016; Chen,. et al. 2017;). In recent study, six membrane proteins exhibited up-regulation in response to heat stress in both varieties of hard fescue. They were categorized into functional groups of photosynthesis (Rubisco activase, disease resistance protein 1, OEE1), stress defense (stromal 70 kDa heat shock-related protein, disease resistance protein 1, CPN 60) and protein degradation (ATP dependent zinc metalloprotease protein) (Wang J et al. 2017). In our study, we found ten differential membrane proteins included light-harvesting Chl a/b-binding (LHC) protein, ATP synthase subunit alpha, ATP synthase subunit beta, photosystem I P700 chlorophyll a apoprotein A2, photosystem II CP43 reaction center protein, photosystem II D2 protein and photosystem II OS from two different tolerant wucai cutivars. The phenomenon of this may because of several proteins were up-related to the same level in both wucai cultivars.

Chlorophyll a-b binding proteins (LHC), which are major components of light-harvesting antennae of PSII, play distinct functions for regulating light-harvesting events, such as the dissipation of excessive light and optimization of light energy utilization (Wang et al. 2015). It has been reported that plants prevent chlorophyll loss from thylakoid membranes by stabilizing photosystem complexes (Shu et al. 2015), such as induces an aggregation of the light-harvesting complex of photosystem II (Tang et al., 2007). Under heat stress or intense light, enhanced amounts of reactive oxygen species will react with proteins and lipids, and will induce various types of photodamage. Therefore, the quality and quantity control of the LHC protein complex is required to avoid photodamage by alleviating excitation energy pressure (Teramoto, H et al. 2002). In detached rice leaves, Kang et al. (2009) concluded that among thylakoid complex proteins, LHCb1, LHCb2, LHCb3 and LHCb5 were not appropriate as senescence-related protein markers due to the stability of LHCII complexes. Our present study have found that WS-1 had more LHCII protein than WS-6 under high temperature stress which was comfirmed by higher ABS/CS_M_, TR_0_/CS_M_, DI_0_/ CS_M_ in WS-1. ATPase are key membrane-bound enzyme complexes for ATP generation, responsible for converting ADP to ATP by using transmembrane proton gradients in the electron transport process in both photosynthesis and respiration (Hopkins, 1999). The ATPase complex consists of α-subunits and β-subunits forming the catalytic core of the enzyme complex with the β subunits involved in catalytic activities and the α subunits being regulatory (Lee et al., 2007). Previous study reported that heat stress could decline the abundance of all a-subunits of ATPase may indicate the impairment of regulatory functions of this enzyme for ATP production under heat stress (Jespersen D et al. 2015). In our present study, both cultivars had a few abundance of the α-subunits of ATPase. While WS-1 maintained a higher abundance of the α-subunits of ATPase than WS-6 could contribute to more active regulatory activity for ATP production under heat stress. Unlike the α-subunits of ATPase, both cultivars had greater abundance of the β-subunits of ATPase in heat stress (Fig. 13). As the β-subunit is the key element for catalytic functions of ATPase, the greater accumulation of β-subunits in WS-1 could facilitate the maintenance of catalytic activities of ATPase for ATP production under heat stress. Several other studies found that ATP synthase was impaired by heat stress (Ferreira et al., 2006; Majoul et al., 2004). The interruption of ATPase function for ATP production is a major culprit of heat stress damages in plants as many processes, including stress defense and repair mechanisms, depend on energy availability for plant survival of long-term heat stress (Crafts-Brandner and Salvucci, 2000).

The reaction center core of PSI contains the special chlorophyll-protein, P700, which is normally surrounded by the Chl a of the core antenna. P700 oxidation is directly linked to the protection of PSI against photoinhibition (Shimakawa, G. et al. 2016). It is reported that the most sensitive components of the photosynthetic apparatus in plants to high temperatures is PSII (Srivastava et al. 1997), whlie in our study, fewer abundance of the P700 proteins in WS-6 may indicated that PSII may influenced by heat stress.

The photosystem II (PSII) reaction center core (RCC) complex of higher plants, algae, and cyanobacteria can be subdivided into a heterodimer containing D1 and D2, the antenna proteins CP47 and CP43, and a large number of low molecular weight integral membrane proteins including the α and β subunits of cytochrome b559 (Yamamoto Y. et al. 2008).

The D1 and D2 proteins form a heterodimer at the center of the PSII complex and serve as the reaction center-binding proteins. They carry essential redox components for charge separation and the subsequent electron transport reaction, such as the reaction-center chlorophyll P680 (the primary electron donor), the primary electron acceptor pheophytin (Pheo), the first and second electron acceptor plastoquinones Q_A_ and Q_B_, the secondary electron donor Tyr_Z_, and the Mn_4-_Ca cluster (Yamamoto Y. et al. 2008). It is reported that the D1 protein had close relationship between nearby proteins such as D2 protein and CP43. Once heat stress occurred, D1 protein damaged is subsequently degraded by proteolytic enzymes or forms specific aggregates with the D2 protein or CP43 (Aro et al. 1993; Yamamoto 2001). In our present study, WS-1 had greater abundance of D2 protein than WS-6 indicated that WS-1 had more complete structure. Meanwhile, WS-1 had high F_V_/F_M_ and PI_ABS_ than WS-6 under high temperature could confirmed.

CP43 is a chlorophyll-protein complex that funnels excitation energy from the main light-harvesting system of photosystem II to the photochemical reaction center (Reppert, M et.al 2008). In our study, ABS/CS_M_ was sharply decreased under high temperature might be due to the fewer abundance of CP43 in WS-6, proved by degredation of antenna proteins (Fig. 13). The identification of proteins regulated by heat stress could lead to a better understanding of the cellular response to dehydration, which is an important and fundamental part of improving the stress tolerance of wucai cultivars.

## Conclusion

According to above data, it was demonstrated that the ability of wucai plants to minimize the heat stress depended upon the growth self-regulation, effectiveness of the photosynthetic and chlorophyll fluorescence, membrane fatty acid composition and protien complexes, differential proteins which varied according to the plant cultivar. Our results revealed that the cultivars WS-1 relatively exhibited heat tolerance of the cultivars. Compared with the heat sensitive cultivars WS-6, WS-1 had a greater capacity for maintaining growth level, photosynthetic parameters and chlorophyll fluorescence. In summary, this study demonstrated that heat tolerance in wucai, as evaluated by membrane stability, as well as leaf photochemical efficiency was associated with greater increase of saturate fatty acids (16:0 and 18:0) content or decrease of unsaturate fatty acids (16:1, 18:1, 18:2 and 18:3) content, as well as less severe down-regulation of membrane proteins and greater up-regulation of heat responsive proteins (Fig. 13). Modification of those membrane constituents could lead to improvement in heat tolerance for wucai and other cool-season vegetable species. Those membrane constituents could also be used as biochemical markers to select for heat-tolerant germplasm due to their contribution to heat tolerance.

## Acknowledgments

No conflict of interest exits in this article, and it is approved by all authors for publication. This work was funded by National Key R & D Program of China (2017YFD0101803), Major Science and Technology Projects of Anhui Province, China (17030701013) and National Natural Science Foundation of China (No. 31701910).

